# Rethinking large scale phylogenomics with EukPhylo v1.0, a flexible toolkit to enable phylogeny-informed data curation and analyses of diverse eukaryotic lineages

**DOI:** 10.1101/2024.08.19.607962

**Authors:** Laura A. Katz, Auden Cote-L’Heureux, Marie Leleu, Godwin Ani, Rebecca Gawron

## Abstract

Eukaryotic diversity is largely microbial, with macroscopic lineages (plant, animals and fungi) nesting among a plethora of diverse protists. Understanding the evolutionary relationships among eukaryotes is rapidly advancing through ‘omics analyses, but phylogenomics are challenging for microeukaryotes, particularly uncultivable lineages, as single-cell sequencing approaches generate a mixture of sequences from hosts, associated microbiomes, and contaminants. Moreover, many analyses of eukaryotic gene families and phylogenies rely on boutique datasets and methods that are challenging for other research groups to replicate. To address these challenges, we present EukPhylo v1.0, a modular, user-friendly pipeline that enables effective data curation through phylogeny-informed contamination removal, estimation of homologous gene families (GFs), and generation of both multisequence alignments and gene trees. Analyses can use a ‘hook’ database of ∼15k ancient GFs or users can easily replace this hook with a set of gene families of interest. We demonstrate the power of EukPhylo, including a suite of stand-alone utilities, through analyses of 500 conserved GFs sampled from 1,000 diverse species of eukaryotes, bacteria and archaea. We show improvements in estimates of the eukaryotic tree of life, recovering clades that are well established in the literature, through successive rounds of curation using the EukPhylo contamination loop. The final trees corroborate numerous hypotheses in the literature (e.g. Opisthokonta, Rhizaria, Amoebozoa) while challenging others (e.g. CRuMs, Obazoa, Diaphoretickes). We believe that the flexibility and transparency of EukPhylo sets standards for curation of ‘omics data for future studies.

**AUTHOR SUMMARY:** The majority of eukaryotic lineages are microbial, with plants and animals nesting among diverse amoeba, flagellates, and other predominantly-microbial clades. Yet analyses of the evolution of microbial eukaryotes is hampered by the lack of tools for efficient analysis of genome-scaled data, especially in light of the genome complexity and high levels of contamination associated with these microorganism. Furthermore, many existing analyses rely on boutique datasets and decision making that lacks transparency and are thus difficult for others to repeat. To address these challenges, we present EukPhylo as an easy-to-use toolkit that fills a gap in phylogenomic methods, and we analyze of 500 gene families from 1,000 species to demonstrate the power of this approach in estimating the eukaryotic tree of life. EukPhylo enables exploration of eukaryotic evolution in a manner that is both transparent and easily repeatable, and hence can be used to illuminate the origin and diversification of eukaryotic life on Earth.

## Introduction

Most of our knowledge about the nature and evolution of eukaryotic life has emerged from studies of macroscopic organisms, with a focus on ‘model’ lineages such as *Drosophila* and *Arabidopsis*. However, such models represent relatively narrow slices of the eukaryotic tree of life (EToL) as the bulk of eukaryotic diversity is microbial [e.g. 1,2]. Insights from microbial eukaryotes (a.k.a. protists) expand our understanding of the ‘rules’ of evolution by their tremendous diversity of morphologies, life cycles and genome properties [3,4]. The gap in knowledge about microbial eukaryotes can be most efficiently filled through taxon-rich phylogenomic analysis methods. However, current practices often rely on boutique datasets and decisions (e.g. in removing contaminants and identifying orthologs) that lack independence and can be challenging to replicate [e.g. 2,5–7]. To address these challenges, we developed EukPhylo v1.0, a flexible phylogenomic pipeline designed for replicable analyses of diverse eukaryotes. EukPhylo includes curated datasets from diverse lineages, a workflow to process omics data and to deploy phylogeny-informed contamination removal, and a suite of utilities to enable efficient estimation of gene families and phylogenies.

Phylogenomic inference faces numerous challenges, including incongruence among loci, long branch attraction [8,9], and lateral gene transfer, which confounds inferences [10]. These issues are especially prevalent in microeukaryotes, where whole genome assemblies are still rare: moreover, many microeukaryotes possess their own microbiome, often resulting in high levels of contamination in transcriptomic samples. Such incongruences lead to conflicting and often spurious tree topologies that can be mitigated by careful selection of taxa and thorough curation of data [11,12]. Another issue that is exacerbated for studies of diverse eukaryotes is frequent reuse of gene families, and even orthologs in concatenated analyses [e.g. 13–16], as this violates the assumption of independence that lies at the heart of phylogenetics [17–19]. The EukPhylo pipeline addresses this non-independence by allowing users to select from our database of ∼15,000 conserved gene families and then to automate ortholog selection for concatenation.

The recent increase in molecular data and bioinformatic methods has spurred the creation of numerous pipelines to infer homology, multisequence alignments (MSAs), gene trees and species-level phylogenies [e.g. 20–23]. These phylogenomic tools differ in their intentions, allowed inputs (e.g. GenBank vs. user-generated data), and intended outputs (e.g. MSAs, trees); yet few include the type of curation needed for analyses of data from microbial eukaryotes given issues with contamination (i.e. from microbiomes and environmental sequences). The first step of many pipelines is to collect homologous sequences, which can be gathered directly from public databases such as GenBank [24], Pfam [25], or OrthoDB [26]. Many recent pipelines rely solely on BLAST [27] or other similarity-searching algorithms (e.g. USearch [28], VSearch [29], and Diamond [30]) to infer homology. However, BLAST is based on similarity only and does not take into account biological relationships [31], and further processing is necessary to confidently establish the source of sequences as well as homology. Phylogenomic pipelines generally include a multi-sequence alignment (MSA) step, which can be challenging when dealing with data from diverse eukaryotes that span ∼1.8 billion years of evolution [32]. For the subsequent estimation of species trees, recent phylogenomic approaches include methods that use gene trees as inputs in inferring species-level relationships [33]. Such methods have been used for projects like the Open Tree of Life [34], and in studies of plants [35], animals [36] and viruses [37]. An example of pipelines that follow these general steps is NovelTree, which performs homology assessment and gene tree construction, though it only accepts protein sequences as input data [20]. Other examples that accept nucleotide sequences are PhyloTa [23], which focuses on homologous identification and collection, and Sumac [22], which focuses on supermatrix building. One pipeline, PhyloFisher, allows users to add new data to a manually curated set of 204 genes that have been used in estimating eukaryotic relationships, but it does not enable *de novo* (aka ‘on the fly’) exploration of contaminants, sequence statistics, or alternative gene families [38].

The importance of a taxon-rich dataset for estimating phylogeny accurately is well established [39,40], and adding diverse lineages (e.g. taxonomic position, rates of evolution, levels of missing data) can improve estimates of species relationships [41–44]. However, even recent estimates of the EToL rely on relatively few taxa (e.g., 234 taxa in Burki et al., [45], 186 taxa in Al Jewari & Baldauf [46], 158 taxa in Cerón-Romero et al., [47] and 109 taxa in Strassert et al., [48]), and many groups now resample the same genes/data matrix in generating species trees [15,e.g. 45,48,49]. The availability of user-friendly tools that facilitate the parallel processing of large numbers of taxa, therefore, has the power to increase the accuracy of large-scale estimates of eukaryotic phylogeny.

Here we present EukPhylo version 1.0 that supports taxon-rich analyses of gene families and gene trees through extensive data curation, and that includes a suite of stand-alone tools plus curated databases. EukPhylo, parts of which are based on a pre-existing pipeline PhyloToL [50,51], includes two main components, which we refer to as EukPhylo parts 1 and 2. EukPhylo part 1 takes input sequences from whole genome or transcriptome assemblies, applies several curation steps, and provides initial homology assessment against a customizable database of reference sequences to assign GFs. EukPhylo part 1 outputs curated coding sequences with gene families assigned, as well as a dataset of descriptive statistics for each input sample. EukPhylo part 2 is highly modular: for a given selection of taxa and GFs, it stringently assesses homology and produces MSAs by iterating the external tool Guidance [52,53]. From MSAs, EukPhylo part 2 builds gene trees, and then includes an innovative workflow for tree topology-based contamination removal.

In addition to presenting the core pipeline, we describe the results of an analysis of 500 conserved GFs from 1,000 taxa, demonstrating how EukPhylo allows users to explore how varying gene sets, taxon sets, or criteria for contamination removal lead to different biological inferences (e.g. differentiating host vs. contaminant material, phylogeny). To this end, we provide a suite of stand-alone tools that describe tree topologies, and demonstrate the effectiveness of our novel tree-based contamination removal methods in improving tree topologies by assessing the monophyly of clades (e.g. ciliates, dinoflagellates, metazoa) supported by robust synapomorphies as well as larger taxa (e.g. Amoebozoa, Archaeplastida, Opisthokonta, SAR).

## Results

### Overview: The pipeline and accompanying scripts

As described in detail below, EukPhylo v1.0 is a flexible and modular pipeline that enables efficient phylogenomic analysis of eukaryotes and includes phylogeny-informed curation of ‘omics data. Compared to PhyloToL [50,51], EukPhylo v1 streamlines the workflow for assigning gene families to data from transcriptomes and genomes (EukPhylo part 1), and expands options for data curation both before and after producing MSAs and gene trees (EukPhylo part 2). All components of the toolkit are written in Python and are available for download on GitHub (https://github.com/Katzlab/EukPhylo) and Zenodo (DOI:10.5281/zenodo.13323372). To supplement and expand the power of the core EukPhylo, we also publish several databases – the Hook reference database for GF assignment, sequences files for 1000 taxa both before and after curation – and a set of utility scripts to facilitate analyses of gene families and gene trees.

EukPhylo is designed to take as input assembled transcripts, genomic CDSs or any sequences with names matching simple criteria (i.e. a 10 digit taxon code plus a unique identifier) as described in the methods (see also Supplementary Text). Curation steps are built into both parts of the pipeline, first enabling analysis of data within a taxon based on sequence properties (e.g. GC content, codon usage; Table S1; File S3) and second using homology assessment and phylogeny-informed removal of contaminants (Fig. 1). EukPhylo part 1 allows users to either use the built-in Hook Database of ∼15,000 eukaryotic gene families (GFs) or a custom reference database to produce curated amino acid and nucleotide sequences for each taxon (Fig. 1). EukPhylo part 2 takes these files as input and constructs an MSA and gene tree for each input gene family (Fig. 1). EukPhylo part 2 also includes a novel workflow for phylogeny-informed contamination removal that we refer to as the ‘contamination loop’, which identifies likely contaminant sequences based on the topology of the gene trees and user-determined criteria, and then removes these sequences, writing them out into a file that users can publish to increase the transparency of their curation methods.

**Fig. 1.**
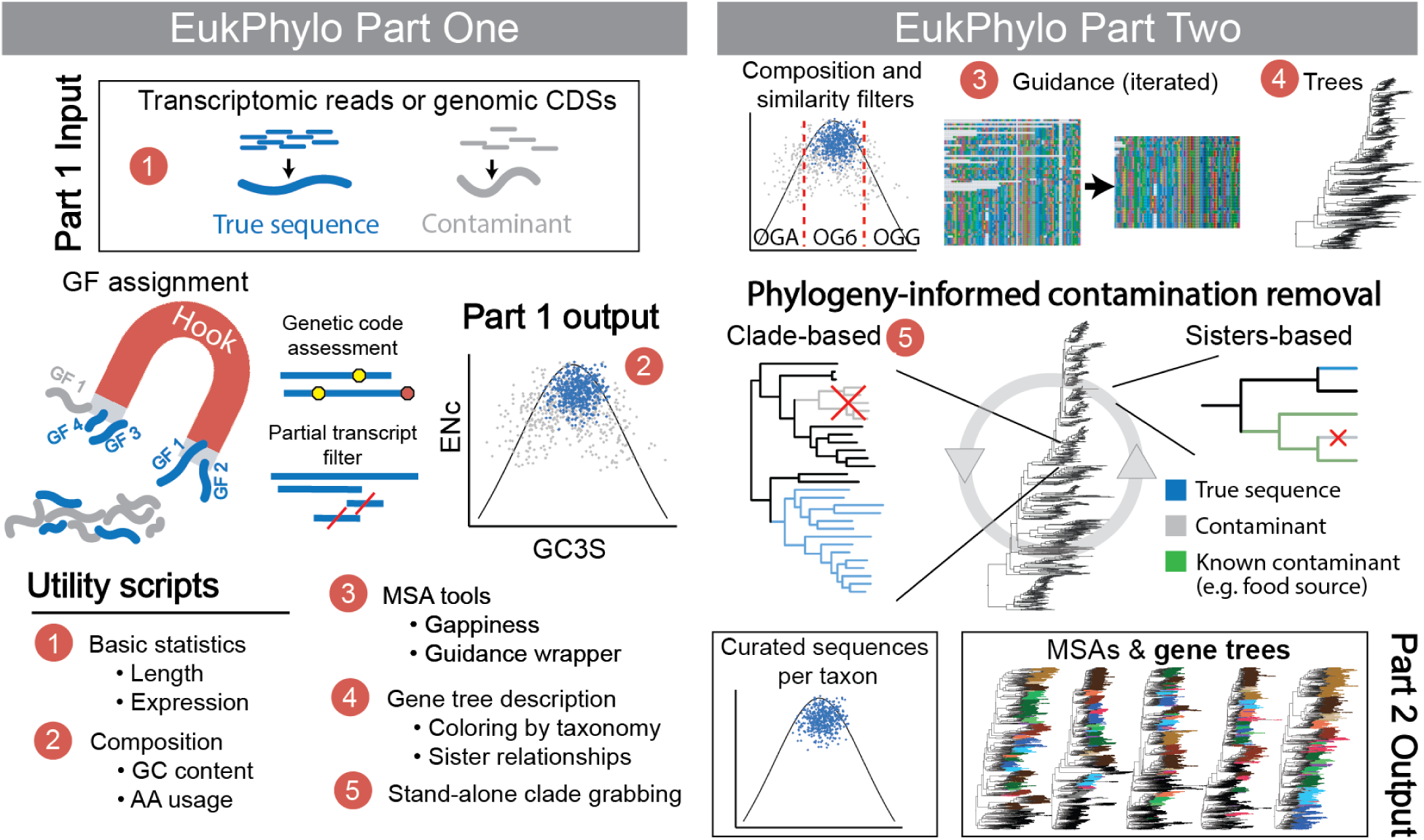
A schematic of the EukPhylo v1.0 core pipeline. EukPhylo comprises two main components, Part One and Part Two. Part One is primarily intended to apply preliminary filtration steps and assign gene families using a reference database. This reference database can be the Hook version 1.0 as described in the main text, or a custom database. Part One takes as input assembled transcripts or genomic CDS, and is able to flexibly handle a variety of genetic codes. In the graph under the “Part 1 Output” heading we show using silent-site GC content (GC3S) vs. the effective number of codons (ENc) that the true sequences from the sequenced sample (blue) tend to have similar composition, with contaminant sequences (gray) lying outside of this range. Part Two builds MSAs by iterating Guidance multiple (by default 5) times for rigorous homology assessment of each gene family, and then builds gene trees. We present a novel phylogeny-informed approach to contamination removal, where contamination is removed from trees in an iterative fashion, either by keeping only sequences in robust clades (“Clade-based”) or removing sequences sister next to known contaminants (“Sisters-based”). We also exemplify the suite of utility tools accompanying this core pipeline, identified by numbers (red circles) where the tool can be applied.

Beside the main pipeline, EukPhylo includes a set of stand-alone utility scripts that aim to increase the power of the analysis done with or without the core EukPhylo pipeline. We divide these scripts into five main categories: basic statistics, composition tools, MSA tools, gene tree description, and contamination removal (Table S2). These tools can be used with EukPhylo output or any other fasta files and/or Newick strings, so long as taxon names have been modified to match the “10-digit” criteria used by EukPhylo.

### Databases

Alongside the suite of scripts that make up the EukPhylo v1.0 toolkit, we provide several taxon-rich databases useful in broad analyses of eukaryotic phylogeny: 1) our Hook reference database for GF assignment, 2) amino acid and nucleotide sequence files for 1000 species with GFs assigned (called ReadyToGo files), and 3) curated MSAs and trees for 500 conserved gene families. The Hook Database is composed of 1,453,081 sequences across 15,414 GFs (Table S3, File S1), and is used in EukPhylo part 1 as a reference database to assign either assembled transcripts or coding domain sequences to GFs. The Hook Database captures a broad diversity of eukaryotic gene families and was built using sequence data from OrthoMCL version 6.13 [54], which we sampled to select for GFs that are present across the eukaryotic tree and/or present in under-sampled lineages of eukaryotes (see methods; Fig. S1, Fig. 2). To add value for users, we also include functional annotations for each GF in the Hook (Table S4; see methods in Supplementary Text). Alternatively, users can insert their own Diamond-formatted database *in lieu* of the Hook, to target only specific genes of interest.

**Fig. 2.**
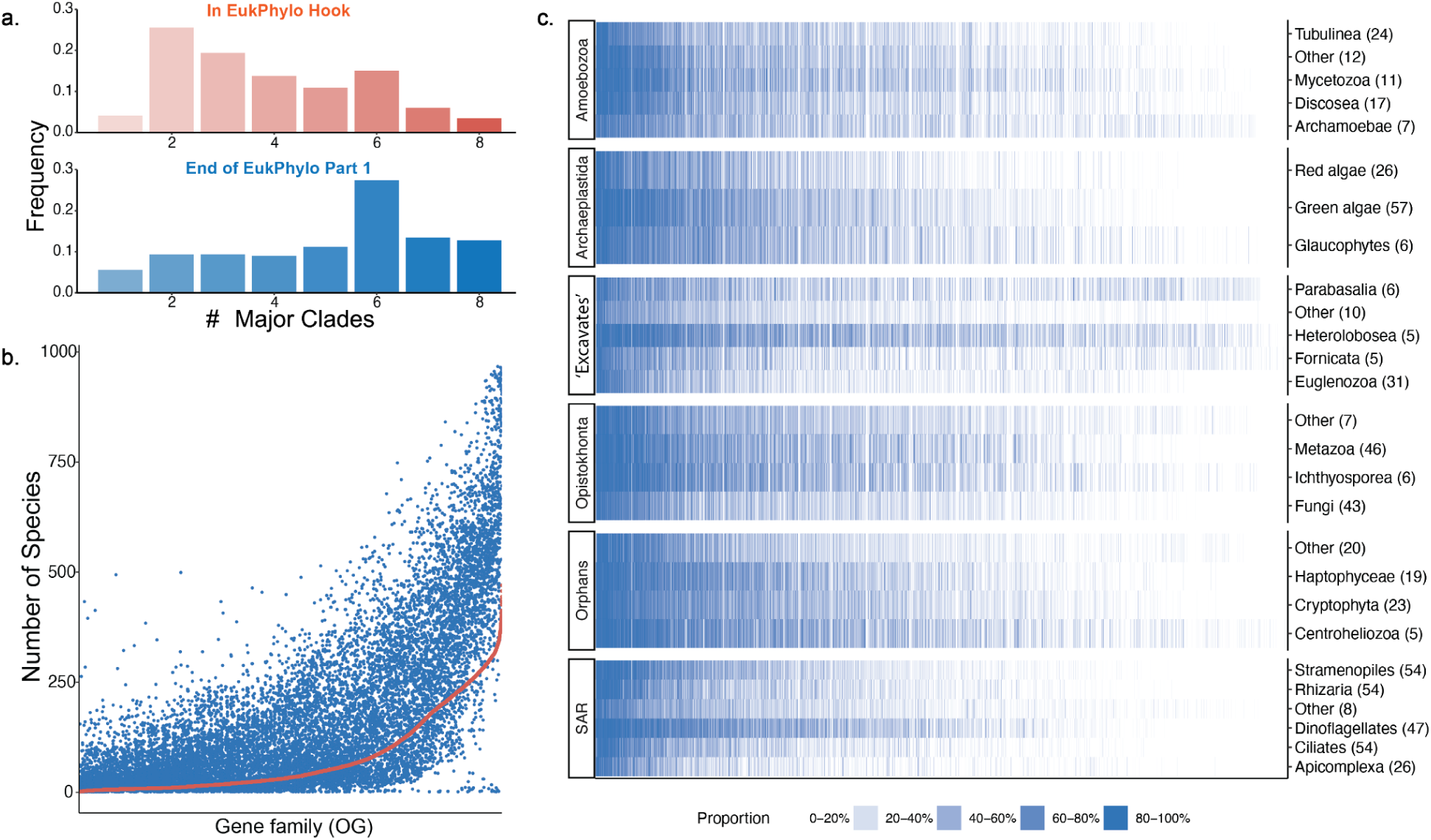
The Hook reference database, which is used in EukPhylo part 1, effectively captures taxonomic diversity in OG assignment. While most OGs as represented in the Hook database are only present in 2-4 major clades, once assigned to our more diverse dataset, most OGs are present in 5-8 major clades, with a mode of 6 (the number of eukaryotic major clades) (a). The number of species with an OG in the ReadyToGo files (blue) correlates with the number of species with that OG in the Hook (red), with very few OGs losing diversity (b). Panel (c) describes the proportion of species (intensity of color) in each eukaryotic minor clade (rows) in which each OG (columns) is found. OGs found in more species are on the left, and those with fewer species are on the right.

### Performance

To assess the performance of the EukPhylo part 1, we compared the speed in processing assembled transcriptomes and genomes through to ReadyToGo files. Using a desktop computer (iMac Pro 2017, 64GB of RAM,10 cores) and a high performance computing cluster (HPC; 128GB of RAM, 24 cores), we processed 10 and 100 transcriptomes and genomes (Table S5). As expected, processing genomes with EukPhylo part 1 was considerably faster compared to the transcriptomes on both computers as coding domains are already called for genomes. On the desktop computer it took roughly 2 hours for 10 transcriptomes (513,904 transcripts) and 24 hours for 100 transcriptomes (3,294,484 transcripts) while the same datasets took 2 and 16 hours respectively on the HPC. Processing the genomes, it took 1 hour 20 minutes and 24 hours on the desktop computer for 10 (106,249 CDS) and 100 taxa (1,158,224 CDS) respectively, and 25 minutes and 21 hours on the HPC to run the same set of taxa. This demonstrates the feasibility of running EukPhylo on desktop or even laptop computers if an HPC is not available.

### Demonstrating the toolkit: Exemplar analysis of 500 conserved gene families

To demonstrate the capabilities of EukPhylo version 1.0, we conducted phylogenomic analyses on 500 gene families using 1000 diverse taxa (275 bacteria, 98 archaea, and 627 eukaryotes (Tables S1,S6-S7 and Files S2-S4). Here our goal is to demonstrate the power of EukPhylo in terms of data curation and in providing replicable approaches to generating both gene and species trees, which includes tracking removed sequences. To this end, we estimate species trees at four different points in the contamination removal process (Fig. 4a-d): 1) before the contamination loop: after generating MSAs/trees from curated “ReadyToGo” files (the output of EukPhylo part 1) based on GC content at silent sites (e.g. GC3S; Table S1 and File S2); 2) after applying sister-based rules to iteratively remove sequences determined to be potential contaminants based on user-established rules (Tables S8 and S9); 3) after clade-grabbing by retaining only sequences for which we have the greatest confidence based on user-established expectations of taxon density (Tables S10 and S11); and 4) after removing gene families that include putative endosymbiotic gene transfers (EGTs), including both primary EGTs (e.g. photosynthetic eukaryotes nested among bacteria) and/or secondary EGTs (e.g. photosynthetic eukaryotic clades interdigitated with one another; File S5). With the goal of demonstrating the power of the approach, we provide readers with details on the presence/absence of clades with robust synapomorphies (e.g. dinoflagellates, ciliates, metazoa, fungi) as well as higher taxonomic groups (e.g. Amoebozoa, Archaeplastida, Opisthokonta, SAR). All files related to this analysis, for each step of the contamination loop, can be found on Figshare (https://doi.org/10.6084/m9.figshare.26527984) in demonstration of another powerful aspect of EukPhylo: the ability to track intermediate files and removed sequences.

### Gene family analysis

We assigned sequences from our 1,000 focal taxa (Table S7) to gene families from our Hook Database using EukPhylo part 1. We selected 500 gene families based on taxonomic presence using two criteria: 1) they are among the most shared GFs in our 1000 taxa and 2) these GFs have relatively low paralogy, making analyses more efficient (Table S13).

Despite the fact that the starting OrthoMCL database is biased in terms of taxonomic availability (e.g. biased towards parasitic lineages [54]), the 15,414 GF Hook Database assigned gene families to a broad diversity of taxa, including poorly represented taxa like *Telonema,* Centrohelidae and other orphan lineages (labeled EE for “everything else”; Fig. 2, Table S1). In fact, the taxonomic distribution of major clades in our ReadyToGo files is greater than in the Hook itself, with more than 75% of the GFs present in at least 4 major clades in the ReadyToGo files (Fig. 2a,b). Nevertheless, the distribution of GFs across taxa is highly variable, reflecting at least two phenomena: the differences between transcriptome and whole genome data, and the prevalence of gene loss in some lineages (e.g. fungi [55] and parasites; Fig. 2c).

Using EukPhylo utilities (CUB.py, GC_identifier.py), we further refined data based on taxon-specific GC content ranges (Supplementary Text) to produce ReadyToGo files with sequences labeled by composition (OG6 if in GC3S range for taxon, OGG and OGA if more GC rich or AT rich, respectively). We provide the resulting ReadyToGo databases for users interested in placing their taxa into a phylogenomic context; users might choose to include all sequences knowing that those labeled OGG or OGA are outside of the GC content ranges chosen for this study (File S2).

### Initial MSAs and gene trees

A first result of EukPhylo part 2 is the generation of 500 MSAs and gene trees that emerge from the curated ReadyToGo data. These 500 MSAs and trees, built after five iterations of Guidance and labeled as “pre-contamination loop,” vary in taxon presence and number of sequences (Table S13), with a total of 15,486 sequences (File S7) removed by Guidance as putative non-homologs. Notably, individual gene trees at this stage include numerous contaminants (e.g. bacterial and food source sequences from transcriptome data, Fig. 3b). To summarize the phylogenetic signal in these data, we generated a concatenated tree (Fig. 4a), which is largely consistent with the published literature; clades that are not recovered in this first analysis (e.g. metazoa, ciliates) are most affected by contamination. For all species tree alignments, we masked columns with ≥95% and ≥50% missing data and we also generated species trees using Asteroid [56] at each step (Fig S3, File S9). The resulting trees from all analyses are discussed in more detail below.

**Fig. 3.**
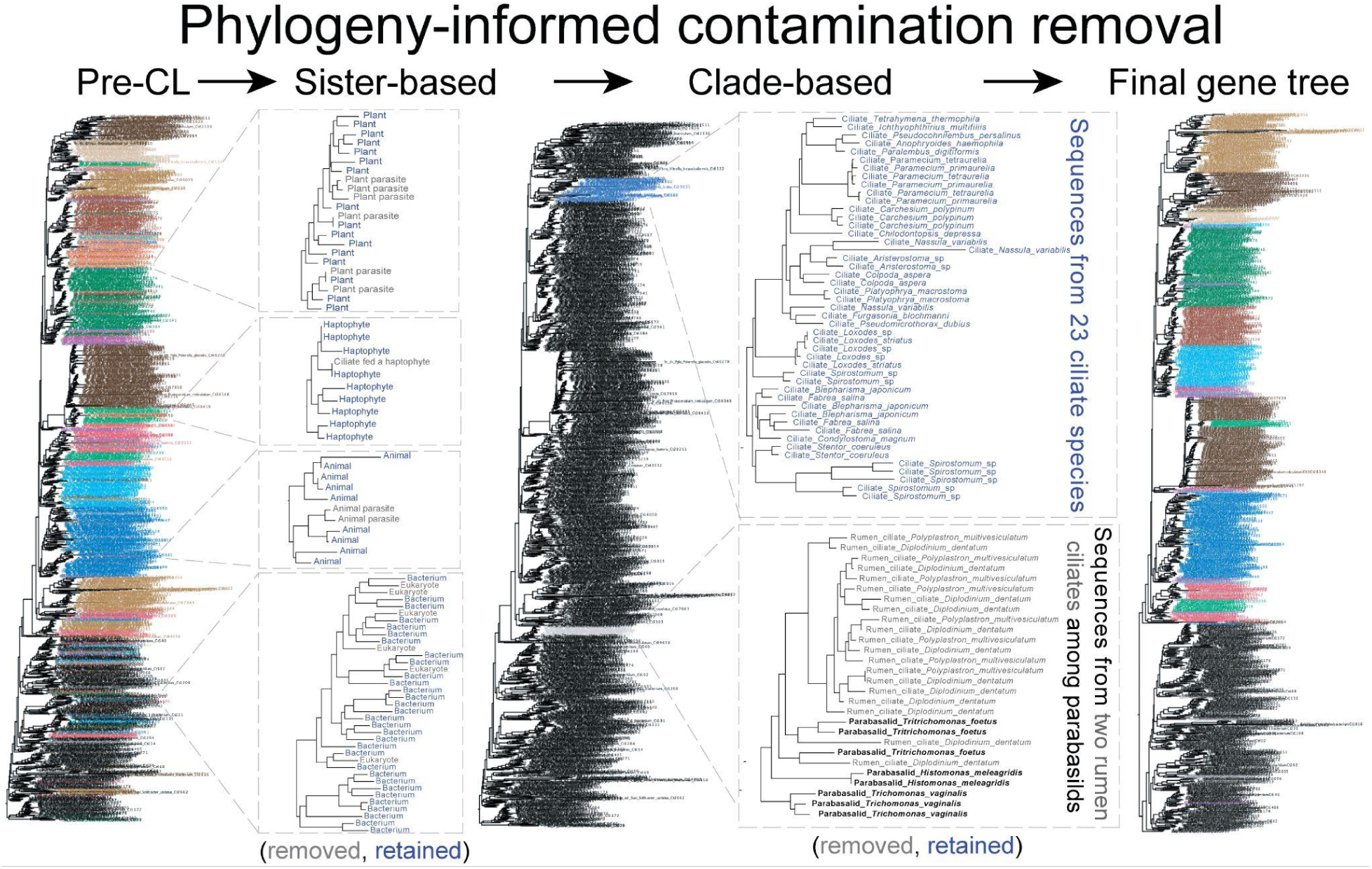
A cartoon depicting phylogeny-informed contamination removal, which is a component of EukPhylo part 2. Users can use the contamination loop to iteratively remove sequences based on their sister species in single-gene trees. Depicted here are sequences (gray) that are either from hosts in analyses of parasites (upper left) or bacterial sequences that come as contaminants in analyses of eukaryotic transcriptomes (lower left). In a second method of contamination removal, users can ‘grab’ (retain) sequences falling in monophyletic clades that meet user-specified robustness criteria (e.g., minimum target group species count and maximum number of non-group species). In the case depicted here, we identified substantial contamination of a subset of ciliate transcriptomes by parabasalids with which the ciliate species are known to share an environment (cow rumen). To remove this ciliate contamination, we used EukPhylo to retain only ciliate sequences falling in clades with at least 12. For clade-based contamination removal, an example of a retained clade is given in blue, and a removed clade in gray in the bottom right.

**Fig. 4:**
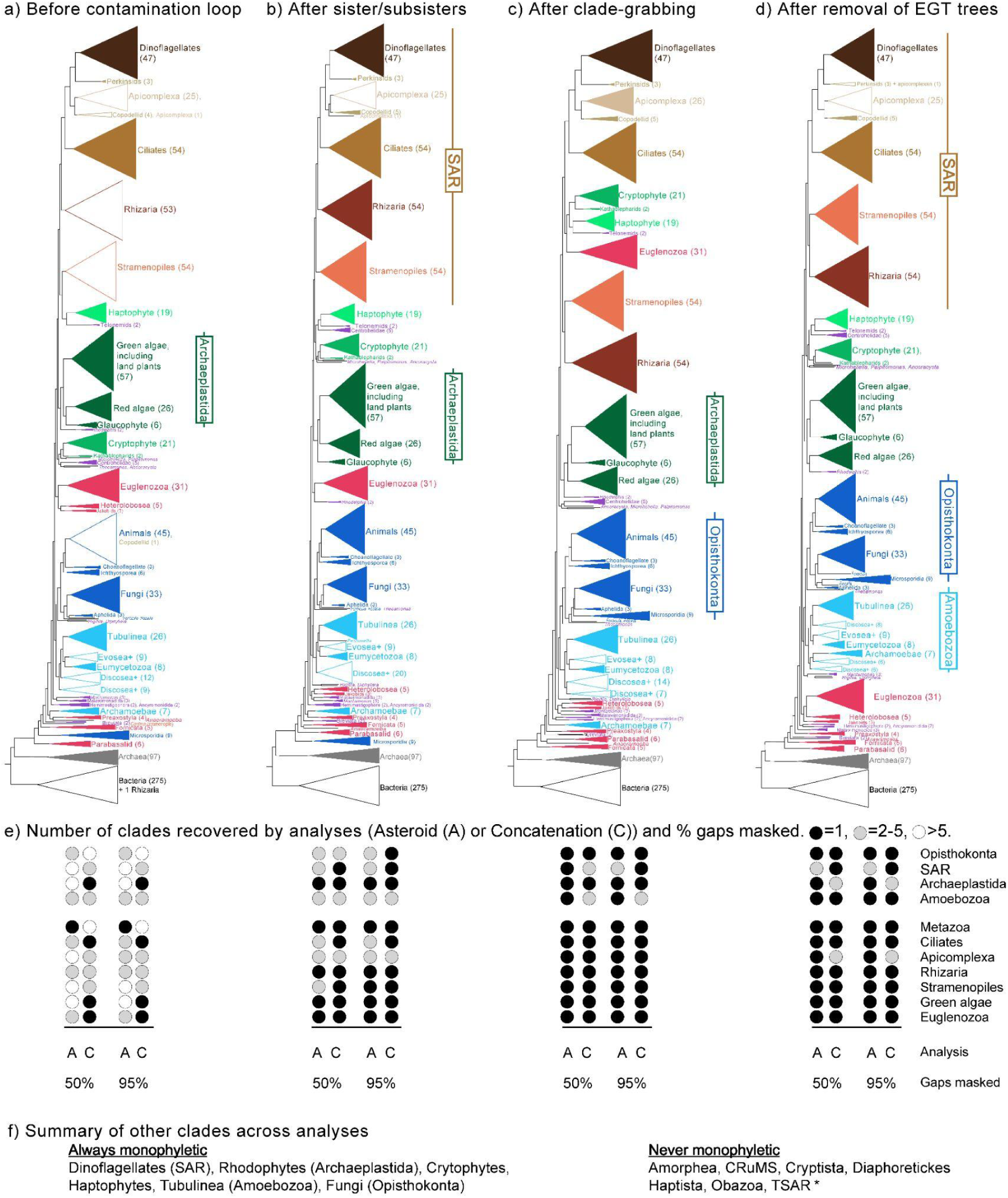
a-d) Concatenated analyses (50% gap trimming) at four stages in the contamination-removal process are generally concordant with published hypotheses as most morphological-defined clades (e.g. dinoflagellates, green algae, Euglenozoa, stramenopiles) are recovered consistently. The four analyses are: a) before contamination loop, b) after removal of contaminants based on sister/subsister rules; c) after clade grabbing to keep ‘best’ sequences; and d) after removal of trees possibly affected by both primary and secondary endosymbiotic gene transfer (EGT). Notably, the monophyly of Opisthokonta emerges after clade grabbing (c) while the monophyly of Amoebozoa and SAR only appear after removing trees affected by EGT (d). Some “orphan” lineages (purple) are stable across trees (i.e. telonemids are always sister to haptophytes, breviates are always towards the root) while other lineages (e.g. Centrohelidae, Hemimastigophora + Ancyromonida) move position across trees; this likely reflects a combination of the effects of missing data and a lack of close relatives. Across each stage of the contamination removal process, the number of key eukaryotic groups that are monophyletic increases in both Asteroid (A) and concatenated (C) analysis under 50% and 95% gap trimming (e). Finally, f) we report groups that are always monophyletic and others that are never found; * indicates that TSAR is recovered only in the Asteroid analysis (50% gap trimmed) after clade-grabbing.

### Contamination Loop

To demonstrate the power of EukPhylo, we ran the “contamination loop” included in EukPhylo part 2 to estimate the topology of the EToL in a manner that is both transparent and repeatable. This contamination loop allows both “sister/subsister” removal (i.e. removal of potential contaminants) and “clade-grabbing” (i.e. retention of sequences for which we have greatest confidence based on taxonomic density) with user-defined rules (see methods; Tables S8 to S11). Using the set of scripts provided with the EukPhylo toolkit (e.g. ContaminationBySisters.py and CountTaxonOccurrence.py, see methods), we assessed taxon presence and sister relationships across initial gene trees to establish three sets of rules: sister, subsister, and clade grabbing (Tables S8 to S11). We make no claim to be correct in all cases, but we believe that EukPhylo’s ability to document sequence choice in a transparent manner is an improvement to best practices in the field.

Examples where sister and subsister rules may be helpful include cases where a taxon, or a pair of taxa, is contaminated by a food source (e.g. ciliate sequences falling among haptophytes), or by a known host in the case of parasites (e.g. the gregarine apicomplexa whose sequences are contaminated by host metazoa; Fig. 3b). We set additional sisters/subsister rules based on our knowledge of the biology of the taxa and inspection of the resulting trees. For example, the Apusozoa *Aphelidium insulamus* and *Aphelidium tribonematis* (Op_ap_Ains and Op_ap_Atri) frequently branch together among stramenopiles or Amoebozoa; we therefore infer that these taxa are likely contaminated by same sources (perhaps in laboratory preparation or in sequencing). The greatest proportion of removed sequences are from taxa within the major clade SAR (abbreviated as Sr), which includes a majority of the field-caught single cell transcriptomic samples (ciliates, foraminifera) from our lab (File S8). At the end of the sister mode of the contamination loop, EukPhylo removed 50,903 of 565,225 sequences (Table S13 and File S8). We again estimated a tree from concatenated alignments masked at 95% and 50% (Fig. 4b) and from Asteroid (File S9), and we discuss the resulting trees below.

The second form of contamination removal in EukPhylo v1.0 is termed “clade-grabbing” as it retains sequences that fall within clades on species trees that meet user-defined criteria. Here, we set minimal numbers of target taxa for clades with many representatives, noting that we likely remove a considerable amount of vertically-inherited data here given the prevalence of gene loss and missing data of some clades. We also identified a list of ‘exceptions’ (i.e. taxa with few close relatives in our analyses; for example orphan lineages as *Mantamonas* and *Hemimastix)* for which all sequences are retained independent of clade size (Tables S10-S12). We first ran clade-grabbing only for ciliates (a clade whose monophyly is not controversial) as we had a strong signal of contamination of parabasalid sequences putative mislabeled as ciliate from species isolated from the digestive system of cows; here, ciliate transcriptomes containing parabasalid sequences (Ex_pa; Fig. 3c) likely cause the ciliates to spuriously fall near parabasalids (Fig. 4b).

After addressing the high level of contamination of ciliate data, we deployed the “clade-grabbing” mode of the contamination loop more generally based on clade sizes determined empirically using the EukPhylo toolkit (CladeSizes.py, see methods). Aiming to retain sequences for which we have greatest confidence, we provide readers with all the information needed to explore removed sequences and/or to implement this procedure for their own data. For example, given that we have a total of 45 diverse metazoa (Tables S1, S6; Op_me; Opisthokonta, Metazoa), we kept only clades containing at least 11 metazoan species. We allowed contaminating sequences to interrupt clades; e.g. for metazoa, we allowed at most 10% of a clade to be non-metazoan species. We share these rules plus the resulting removed sequences in Tables S8-S12 and File S8. In the end, clade grabbing removed 129,458 of 514,272 sequences (Table S13 and File S8). Here again, we emphasize on the importance of transparency and user defined rules for clade grabbing, as this process will most likely remove “good” sequences as well as contaminants, while selecting for the strongest signals in tree topologies.

Given the many papers demonstrating a substantial effect of endosymbiotic gene transfer (EGT) from plastid genomes to nuclear genomes [reviewed in 57], we conducted a final analysis to remove gene families that may have been affected by both primary and secondary EGT. To this end, we used the utility scripts CladeGrabbing.py and CladeSize.py to identify gene-families with a putative photosynthetic history; here we define gene families affected by primary EGT as gene families with photosynthetic lineages nested among bacteria (often cyanobacteria, but allowing other bacterial sisters given the prevalence of gene loss and LGT among bacteria) while secondary EGTs are identified as gene families with the greatest proportion of sequences of intermingled photosynthetic lineages (e.g. dinoflagellates nested among diatoms; Tables S12 and File S5). After removing 113 and 56 GFs possibly affected by primary and secondary EGT, respectively, we ran concatenated and asteroid analyses and we compare the resulting “EGT removal” trees below (Fig. 4d).

### Estimating EToL through iterated data curation

We discuss the topology of EToL from four stages of analyses to demonstrate the power of the tree-based contamination removal workflow in EukPhylo. As a measure of robustness, we focus on the presence of clades whose monophyly is supported through ultrastructure and/or by robust synapomorphies (e.g. ciliates, dinoflagellates, metazoa, fungi, green algae), as well as higher-level taxa (e.g. Amoebozoa, Archaeplastida, Opisthokonta, SAR). We find that tree topologies estimated by both concatenation and Asteroid (Figure S3, Table S12, File S9) are overall concordant through the various stages of curation for the 1,000 taxa and 500 gene families, though with marked improvements through data curation (Fig. 4a-e, with monophyletic clades represented by filled triangles (a-d) or circles (e)). We also provide readers with individual alignments (Files S5-6), individual gene trees (File S9), and lists of removed sequences (File S7-8).

The tree-based contamination removal in EukPhylo enables improvement of tree topologies through transparent removal of putative contaminants. Prior to using the tree-based contamination tools, our taxon-rich concatenated analyses recover many clades with robust synapomorphies across all analyses (50% and 95% gap trimmed, concatenated and asteroid, including fungi, dinoflagellates, cryptophytes, haptophytes, and Tubulinea (Fig 4a; Table S12)). However, the monophyly of some clades is disrupted by single taxa: the monophyly of Rhizaria is disrupted by the placement of the foraminifera *Notodendrodes hyalinosphaira* (Sr_rh_ArpA) among bacteria, and the parasite *Piridium sociabile* (Sr_co_Psoc) falls within animals in concatenated analyses (Fig. 4a, Table S12). Other aberrant observations include the placement of Microsporidia (a lineage known to have elevated rates of evolution [58,59]) and Archamoebae towards the base of the eukaryotic portion of the tree, and the non-monophyly of stramenopiles. Asteroid analyses of these data reveals further evidence of contamination as numerous clades (e.g. ciliates, Euglenozoa, green algae; Table S12) are non-monophyletic, which highlights the impact of contamination in omics data from microeukaryotes.

The iterative contamination removal in EukPhylo improves tree topology as we remove contamination based on user-set ‘sister/subsister’ rules (Tables S8-S9) and then retain only the most robust sequences through ‘clade grabbing.’ Deploying rules for sister-based contamination removal improves the topology of EToL in that we consistently recover clades like metazoa, Euglenozoa, colpodellids and Rhizaria (Figs 4b,e, Table S12). However, the monophyly of ciliates, another clade with robust synapomorphies (cilia and dimorphic nuclei [1]) emerges only after deploying the second tree-based contamination method that we call ‘clade-grabbing’; here, EukPhylo allows users to retain clades with a pre-set number of target taxa (Table S10, File S5), which allowed us to distinguish ciliate signal from contamination by parabasalids among a subset of ciliates isolated from the rumen of cows (see above). Our final curation step, ‘EGT removal’, excluded gene families that may be affected by primary and secondary EGT (Fig 4d, Table S12, File S5, S6). Intriguingly, Archaeplastida (present across most other analyses; Fig 4e, Table S12) are not monophyletic in concatenated trees after ‘EGT removal’ as two members of the genus *Rhodelphis* fall sister to red algae, consistent with previous analysis of these ‘orphan’ species [60].

We also assessed changes in higher-level eukaryotic taxa throughout stages of contamination removal. Opisthokonta (animals, fungi and their microbial relatives) emerges consistently only after sister/subsisters removal for both Asteroid and concatenated analyses (Fig. 4, Table S12). The monophyly of SAR (stramenopiles, alveolates and Rhizaria) and Amoebozoa are recovered in a subset of analyses following contamination removal (Fig 4b-e; Table S12). The orphan lineage Hemimastigophora is consistently sister to the Ancyromonidida, falling nested among ‘excavate’ and orphan lineages towards the root of our trees. The placement of other orphan lineages (purple branches, Fig. 4a-4d; Figure S3, File S9) varies across analyses, with some lineages like Breviata, Malawimonadida and *Mantomonas* falling towards the root of EToL (Fig 4a-d, Figure S3, File S9)), though missing data likely confounds the placements of all of these lineages (see below). Other proposed eukaryotic “supergroups” (e.g. CRuMs, Obazoa, Diaphoretickes) are not recovered in any analysis, and the proposed clade ‘TSAR’ (*Telonema* + SAR) is recovered only in the clade-grabbed trees analyzed by Asteroid with 50% gap-trimming (Table S12, Figure S3, File S9).

Across all analyses, we obtain a root among excavate taxa (formerly the clade ‘Excavata’) plus a few orphan species, though this may be due in part to a high amount of missing data in these lineages. Clades with the greatest proportion of gaps and fewest numbers of gene families, such as Breviata, Fornicata, Jakobida, Malawimonadida, Microsporidia, and Preaxostyla, tend to be most unstable across analyses and to fall close to the root of the tree (Fig 4, Table S14, Fig S4).

## Discussion

EukPhylo v1.0 provides a platform for the efficient curation and analysis of ‘omics data from eukaryotes, using tree-based methods that enable exploration of both gene families and species relationships. Key aspects of EukPhylo are its repeatability, flexibility and transparency, as users can record parameters (e.g. in identifying contaminants) and report both retained and removed sequences through every step. Analyses of diverse microbial eukaryotes, and particularly uncultivable lineages characterized by single-cell ‘omics, require curation to choose most shared gene families, identify homologs and remove contamination (e.g. from contaminants and/or symbionts). Numerous ‘boutique’ approaches that require hand curation have been used to estimate eukaryotic phylogeny [e.g. 13,48,61–63], but reliance on resampling of data (i.e. choosing orthologs to match previous concatenated gene sets) can lead to issues arising from a lack of independence [reviewed in 64]. While standards of curation and data quality have been developed for analyses such as genome assembly and annotation [e.g. 65,66], analogous standards do not yet exist for phylogenomics and we believe that EukPhylo will in part fill this gap by providing transparently repeatable methods.

EukPhylo provides a streamlined method to include diverse data types (e.g. transcriptomes, genomes) allowing users to choose gene families for each study that maximize power in analyses and to curate data *via* both per-taxon and tree-based comparative approaches (Fig. 3). The databases and scripts require minimal effort for installation and are structured for users with modest bioinformatic skills, as accessibility and usability are key considerations in designing scientific tools [67]. We provide EukPhylo with default settings for parameters that we believe are a reasonable starting place for analyses, though all parameters are easily customizable. For those interested in a a taxon-rich dataset for analyzing data from previously uncharacterized taxa, we provide the EukPhylo Database, a set of curated data from 1,000 diverse transcriptomes and genomes from eukaryotes, bacteria, and archaea (Table S1, File S2); these data will enable large-scale phylogenomic analyses and robust inferences about eukaryotic features. Alongside the code, we provide a comprehensive manual on GitHub that describes how to use the EukPhylo toolkit (https://github.com/Katzlab/EukPhylo-6/wiki).

The importance of EukPhylo’s flexibility lies in its ability to help users describe the effect of differing methodologies on biological inferences. EukPhylo allows the exploration of evolutionary questions with reproducibility and transparency. By starting with a taxon-rich dataset and over 15,000 GFs in our Hook Database, we enable users to assess the impact of contamination on their inferences, an improvement over previous phylogenomic literature that relies on reuse of previously-curated gene families. Users can also easily replace the Hook Database with a set of gene families of interest, a step that extends beyond previous phylogenomic approaches [2,5,6]. In addition to varying the input data, EukPhylo users have a large amount of leeway in deciding how to remove putative contamination from their dataset (e.g. by setting rules for sister/subsister, and exploring different numbers of taxa in parameterizing ‘clade grabbing’). Further, as we demonstrate in our EGT analyses, EukPhylo’s suite of stand-alone utility tools allows users to easily compare the monophyly of well-established clades (e.g., ciliates or metazoa) across trees subject to different criteria to explore hypotheses given their particular questions and data.

Our exemplary analysis of 500 conserved gene families demonstrates the power of EukPhylo to analyze large, diverse eukaryotic datasets, and to improve topologies through tree-based contamination removal. Even the first trees (concatenated and asteroid) to emerge from EukPhylo are largely concordant with published literature (Fig. 4a, Table S12), particularly for morphologically-defined clades like dinoflagellates, animals, red algae, Tubulinea, and Euglenozoa [e.g. 1,2,46–48]. Importantly, most other published analyses rely on many fewer taxa and fail to demonstrate monophyly of so many clades. Also, all of our analyses place the root among parabasalids and Fornicata, generally consistent with the hypothesis in Al Jewari and Baldauf (2023), though testing the root would require more attention to gene-family selection and to mitigating the effect of missing data (Fig S4, Table S14). As we remove putative contaminants based on sister/subsister analysis (Fig. 4b) and then retain sequences for which we have the greatest confidence through ‘clade-grabbing’ (Fig. 4c) we see additional major clades supported (e.g. Opisthokonta, Alveolata (ciliates, dinoflagellates, Apicomplexa and their relatives)). Only with removal of gene families most affected by putative EGT do we recover SAR (stramenopiles, alveolates, Rhizaria), and we note the interesting shift in relationships among Archaeplastida in these analyses (Fig. 4d); these analyses suggest that EGT may be a driver in inferences about EToL. Finally, we do not recover a number of eukaryotic ‘supergroups’ like Amorphea, CrumS, Cryptista, Diaphoretickes, Haptista, Obazoa and TSAR (Fig. 4, Table S12), suggesting the possibility that they emerged through resampling of the same data across analyses.

EukPhylo allows for ‘phylogeny on the fly’ as users can reset gene families, contamination-removal rules and run the pipeline and associated toolkit with great flexibility, modularity and transparency. We have demonstrated how EukPhylo can be used as a starting place for testing evolutionary hypotheses. EukPhylo allows researchers to rapidly compare hypotheses regarding the placement of disputed lineages (e.g. Telonemia [48] or Hemimastigophora [14]) in a repeatable and transparent manner (i.e. tracking of removed sequences), while enabling taxon-rich analyses. EukPhylo also allows exploration of the placement of ‘orphan’ lineages on the eukaryotic tree by leveraging the ability to treat data from these taxa differently (e.g. more leniency in curation) than data from taxa belonging to better sampled clades. Moreover, because researchers can choose gene families independently for each study for up to 1000 taxa provided by this study, EukPhylo may help to mitigate the problem of recovering similar topologies across resampled datasets [15,e.g. 45,48,49]. In sum, EukPhylo synthesizes a broad set of tools to facilitate large phylogenomic analyses from start to finish, and most importantly sets foundations for establishing best practices in the field.

There are several important caveats to consider when using EukPhylo. While the EukPhylo pipeline is built to be generalizable, it includes stringent data-quality filters that may remove sequences of interest to certain studies, and is therefore best suited for processing data for large-scale evolutionary or population-level analyses (e.g. generating many diverse gene trees for a supertree approach to phylogeny). Similarly, some parameters which we applied universally (such as the Guidance sequence cut off) are likely not appropriate for all taxonomic groups, and there is room for improving the flexibility of parameter fitting by taxon. Users interested in the evolutionary history of specific genes should use EukPhylo with caution as fast-evolving sequences may be removed, though again assessment here is easy as EukPhylo provides intermediate files as output for inspection by users. Finally, we note that though the stochasticity associated with aligning sequences and building gene trees makes some aspects of analyses not completely replicable, the structure of EukPhylo increases transparency (i.e. by retaining user-defined rules and removed sequences) to enable streamlined and large-scale phylogenomic studies.

### Synthesis

Currently, studies of microbial eukaryotes rely heavily on bioinformatics tools developed for macrobes and/or bacteria; however, such tools do not incorporate workflows that are critical for accurate analysis of eukaryotic lineages where the underlying data must be rigorously cleaned in light of contamination and non-vertical gene transfer. In light of increased attention to the importance of democratizing biology research, especially in the realm of software tools [67–69], we designed EukPhylo to be accessible to researchers with limited bioinformatic background. In its novel phylogeny-informed contamination removal methodology and especially in its flexibility in curating well-sampled lineages differently from ‘orphan’ eukaryotic lineages, EukPhylo has the potential to increase the standards and repeatability of studies of eukaryotic phylogeny.

## Materials and Methods

Here we provide an overview of methods, including descriptions of taxa and gene families, the development of the EukPhylo Hook Database, brief descriptions of the functionality of EukPhylo, and details on our exemplary analyses of 500 gene families in 1000 taxa. Further details are provided in the supplemental methods section of the Supplementary Materials.

### Taxon and gene family designations

EukPhylo v1.0 is based on carefully controlled names of both clades and species that facilitate analyses. Each transcriptome and genome in the EukPhylo database is identified using a ten-digit code, which represents either an individual cell or GenBank accession, or a pool of transcripts as noted in Table S1. The first two digits of the code identify one of eight ‘major’ clades as follows: Ba, Bacteria; Za, Archaea; Op, Opisthokonta; Am, Amoebozoa; Ex, excavate lineages (formerly the clade ‘Excavata’); Sr, SAR (Stramenopila, Alveolata, and Rhizaria); Pl, Archaeplastida; EE, orphan lineages. The next two digits identify the taxonomy of the taxon at the ‘minor’ clade level (e.g. within Opisthokonta are the minor clades Op_me for Metazoa; Op_fu for Fungi; Op_ch for choanoflagellates; and Op_ic for Ichthyosporea; Table S6). The last four digits identify the species and, if applicable, sample ID within a species (e.g. Am_tu indicates the minor clade Tubulinea, and there are multiple samples of *Hyalosphenia papilio*, identified as Am_tu_Hp01, Am_tu_Hp02, etc.; Table S1). Gene families (GFs) are identified as per the notation in OrthoMCL version 6.13 [54], with the prefix OG6_ followed by a unique six number sequence (see sections on Hook Database and composition-based curation below). All sequence identifiers used in EukPhylo databases are unique and begin with the ten-digit taxon identifier, then are labeled by a unique contig/CDS ID designated either by an assembler or by annotations as downloaded from GenBank, and end with a ten-digit GF identifier.

### Development of the Hook Database

As a starting place for evolutionary analyses of lineages sampled across the EToL, we developed a Hook Database of 15,414 GFs selected for presence across a representative set of eukaryotes. The Hook allows assignment of sequences to GFs and can easily be replaced by researchers interested in specific gene families (*e.g.* gene families involved in epigenetics, in meiosis, etc.). To develop the Hook Database (Fig. S1), we started with ‘core’ orthologs from the OrthoMCL version 6.13 database (495,339 GFs). We then proceeded to several curation steps, with the goals of 1) reducing the size of the database as much as possible while retaining diversity within eukaryotes; 2) retaining only GFs that are present in a representative set of eukaryotes given our focus on microbial lineages (i.e. we undersample animal-specific and plant-specific GFs); and 3) removing GFs and sequences within GFs that are likely to cause sequences to be misassigned or assigned to groups of sequences without useful functional meaning (e.g. sequences that comprise only a single common domain, or chimeric sequences). To achieve these goals, we assessed the taxonomic diversity and the quality of each GF using a variety of custom scripts (DOI:10.5281/zenodo.13323372). We detail these curation steps in the Supplementary Materials, and reiterate the goal of generating a set of representative gene families to use in analyses of diverse eukaryotes.

### EukPhylo Part 1

EukPhylo comprises two components: the first (EukPhylo part 1) provides initial gene family assignment to sequences and and the second (EukPhylo part 2) builds alignments and phylogenetic trees. Central to all of EukPhylo is the use of consistent taxon codes (see above). EukPhylo part 1 has two versions, the first intended for use with transcriptomic data, and accepts as input assembled transcripts as produced by rnaSpades (Fig. S2). Users may use other assembly tools as long as sequence names are subsequently formatted in the style of rnaSpades (i.e. including a contig identifier, k-mer coverage and length). The second version is for use with whole genome data, and accepts as input nucleotide coding sequences (CDS). Each step can be run individually across any number of input samples, runs can be paused and resumed at any stage, and this can be flexibly managed using a wrapper script provided in the Zenodo repository (DOI:10.5281/zenodo.13323372)

### Transcriptomic pipeline

The transcriptomic pipeline requires three inputs: a fasta file of correctly named contigs (see manual), a file specifying a genetic code (if known) for each taxon, and for those interested in removing sequences misidentified due to index-hopping, a list of names of conspecifics (i.e. taxa/samples that are expected to share identical nucleotide sequences). As described in detail in the Supplementary Text and in Fig. S2, EukPhylo part 1 removes sequences based on length parameters, and optionally sequences that are likely incorrectly labeled due to index hopping [70,71] in the same sequencing run. Next, putative rRNA sequences are moved to a separate folder and remaining sequences are labeled as possible prokaryotic contamination (ending in _P) for users to inspect downstream. To provide initial gene family assignments, Diamond [54] is used to compare sequences either to the EukPhylo Hook Database (described above) or a user-provided database. As the Hook Database is replaceable and customizable, this step offers an opportunity to filter transcriptomic data for a group of gene families/functional groups of interest. EukPhylo then captures ORFs as both nucleotide and amino acid sequences. Finally, EukPhylo part 1 removes putative chimeric and partial transcripts to produce “ReadyToGo” fasta files and calculates a variety of statistics for both sequences and taxa.

### Genomic pipeline

The version of EukPhylo part 1 applicable to coding domain sequences (CDSs) from whole genome assemblies is similar to the version for transcriptomic data described above and in the Supplementary Text, but with some important differences. Given that coding domains are already determined, this version of EukPhylo part 1 has no length filter, and instead immediately evaluates in-frame stop codon usage and translates the nucleotide CDSs to amino acids, at which point it uses Diamond BLASTp to assign gene families against the same reference database (in our analyses, the Hook Database). Next, the pipeline filters sequences by relative length, removing any sequence less than one third or more than 1.5 times the average the length of its gene family in the Hook Database. After some reformatting, EukPhylo part 1 then outputs the same “ReadyToGo” files as the transcriptome version of the pipeline: a nucleotide and amino acid fasta file with gene families assigned for each taxon, a tab-separated file of BLASTp data against the Hook Database, and summary statistics.

### EukPhylo Part Two

The second major component of the pipeline (EukPhylo part 2) starts from the “ReadyToGo” files produced by part 1 (or any set of per-taxon sequences with names that match PLT6 criteria) and generates multisequence alignments and trees. Prior to running Guidance [52,53] for homology assessment, optional filters are available in the script ‘preguidance.py’, to select the sequences to use for the analysis based on GC composition or high similarity proportions (details in the Supplementary Text), on the whole dataset or on specific taxon.

Then EukPhylo part 2 runs Guidance [52,53] in an iterative fashion to remove non-homologous sequences defined as those that fall below the sequence score cutoff. (We note that there is some stochasticity here given the iteration of alignments built into the method.) After inspecting a diversity of gene families, we have lowered the default sequence score cutoff from 0.6 to 0.3, though this may not be appropriate for all genes (see caveats section below). To remove regions with large gaps that can confound tree building, the resulting MSAs are then run through TrimAl [72] to remove all sites in the alignment that are at least 95% gaps (again, a parameter a user could alter). The last step of EukPhylo part 2 before phylogeny-based contamination removal is to construct gene trees, though users can stop EukPhylo after Guidance to build trees with other softwares as they prefer. Currently EukPhylo supports RAxML [73], IQ-Tree (with the hardcoded protein LG+G model [74], and FastTree [75].

### Phylogeny-based contamination removal

A key innovation in EukPhylo v1.0 is the “contamination loop”, an iterative tool to identify and remove contamination based on analyses of single gene trees. This tool incorporates two main methods of contamination assessment informed by tree topology. The first method – ‘sisters’ mode – is intended to target specific instances of contamination. It enables users to remove sequences based on cases of repeated contamination in target taxa, determined by prior assessment of trees (aided by the utility script ContaminationBySisters.py or known contaminants; Fig. 3). We provide additional details in the Supplementary Materials. The second method – “clade-based contamination removal” – is intended for cases when the user is interested in genes present in a group of organisms with multiple representative samples and/or species in the gene trees (Fig. 3). For a given set of target taxa, this method identifies robust, monophyletic clades containing those taxa within each gene tree (allowing a user set number of contaminants), and re-aligns and re-builds the tree excluding all sequences from the target taxa that do not fall into these robust clades. In both cases, sister and clade grabbing, a user-defined set of rules is necessary and can be built using the set of utility scripts provided with the main pipeline. Given that these methods incorporate tree-building on each iteration, users should expect some amount of stochasticity in which sequences are removed.

### Ortholog selection for concatenation

EukPhylo part 2 includes an option to concatenate representative sequences per GF into a supermatrix from which users can construct a species tree. This can be done as part of an end-to-end EukPhylo run, or by inputting already complete alignments and gene trees and running only the concatenation step. If a GF has more than one sequence from a taxon, EukPhylo keeps only the sequences falling in the monophyletic clade in the tree that contains the greatest number of species of the taxon’s clade as determined by its sample identifier. If multiple sequences from the taxon fall into this largest clade, then the sequence with the highest ‘score’ (defined as length times k-mer coverage for transcriptomic data with k-mer coverage in the sequence ID as formatted by rnaSpades, and otherwise just length) is kept for the concatenated alignment. If a GF is not present as a taxon, its missing data are filled in with gaps in the concatenated alignment. Along with the concatenated alignment, this part of the pipeline outputs individual alignments with orthologs selected (and re-aligned with MAFFT), in case a user wants to construct a model-partitioned or other specialized kind of species tree.

### Conserved OG analysis

To demonstrate the power of EukPhylo, we conducted a phylogenetic analysis on 500 conserved gene families among 1,000 species. Selection of taxa and gene families to include in this study was based on quality of data and taxon presence. We went through several rounds of curation and selection that are detailed in the Supplementary Text; the final selection of taxa is described in Tables S1 and S7. We used EukPhylo part 1 to produce fasta formatted CDS files (genomes) and assembled transcripts (transcriptomes) for each genomes and transcriptomes downloaded from public databases plus data generated in our lab.

We then reran EukPhylo part 2 with these 1000 taxa, using only the sequences labeled as ‘OG6’, based on GC composition (see Supplementary Text for details), with five iterations of Guidance, and built trees using IQ-Tree (-m LG+G; File S4). We then removed sequences identified as contaminants by the contamination loop in EukPhylo part 2. We first ran ten iterations in ‘sisters’ mode, using the rules file provided in Table S8, followed by five iterations of ‘subsisters’ rules on a select number of taxa (Table S9). Next, we ran two separate iterations of the ‘clade’ mode, the first one to remove only the ciliate parasites of Parabasalids (Ex_pa) that occurred when transcriptome data were generated from co-contaminated rumen ciliates, and the second one to remove sequences from all other well-sampled taxa (see Tables S10-S11 for rules, and Supplementary Text for details).

For the final analyses, we removed gene families that showed evidence of either primary or secondary endosymbiotic gene transfer (EGT). We first used the utility script CladeSizes.py to identify trees where multiple photosynthetic lineages nest in a single clade. We identified putative primary EGT events as clades comprising only photosynthetic eukaryotes and bacteria (and occasionally archaea), with many of these including cyanobacteria; we used this broad approach in light of the possibility of either LGTs among prokaryotes (i.e. from cyanobacteria to other prokaryotes) after transfer to eukaryotes, and because of the possibility of multiple sources of photosynthetic machinery in eukaryotes [e.g. 76]. We identified putative secondary (or tertiary) EGT events as cases in which we found interdigitation of multiple lineages of photosynthetic eukaryotes (e.g., photosynthetic stramenopiles nested in red algae). We manually examined all trees with a large number of putative primary and/or secondary EGT events (identified using the utility script CladeSize.py), resulting in a set of 169 OGs total that we removed to construct our final EGT-removed species tree (Fig. 4d).

To build species trees, we used two methods: Asteroid [56] and the concatenation option included in EukPhylo. At each step of the process, we selected orthologs (i.e., removed putative paralogs) and built a concatenated alignment using the methods built into EukPhylo part 2 (see supplemental methods and the EukPhylo v1.0 GitHub wiki page for more information); species trees were then built with IqTREE (-m LG+G, Files S4-S6). We also used Asteroid [56] to build super trees with trees generated by EukPhylo, at each step of the contamination loop (File S9).

## Data and Software Availability

The main EukPhylo pipeline and accompanying scripts, including all scripts used for this study, are available on GitHub (https://github.com/Katzlab/EukPhylo) and Zenodo (DOI:10.5281/zenodo.13323372). All results and outputs generated by this study, including Tables 1 to 15 and Files 1 to 10 listed in the manuscript, are available on Figshare (https://figshare.com/projects/EukPhylo_Supplemental_Files/196552).

## Acknowledgements

We are grateful to Elinor Sterner, Julian Hernandez (Smith College) and Xyrus Maurer-Alcalá (AMNH) for their help in early stages of this study, and to the Unity Cluster (Massachusetts Green High Performance Computing Center) for HPC resources. Recent support for LAK has come from two NSF awards (DEB-2230391, OCE-1924570) as well as funding from the NIH (R15HG01040).

## Notes

https://doi.org/10.6084/m9.figshare.26540599

### Competing Interest Statement

The authors have declared no competing interest.

### Summary of Updates

Changing the name of the pipeline from PhyloToL version 6 to EukPhylo version 1 to reflect the magnitude of the update.

