## Supplementary figures and images for "Rethinking large scale phylogenomics with EukPhylo v1.0, a flexible toolkit to enable phylogeny-informed data curation and analyses of diverse eukaryotic lineages"

### Fig S1

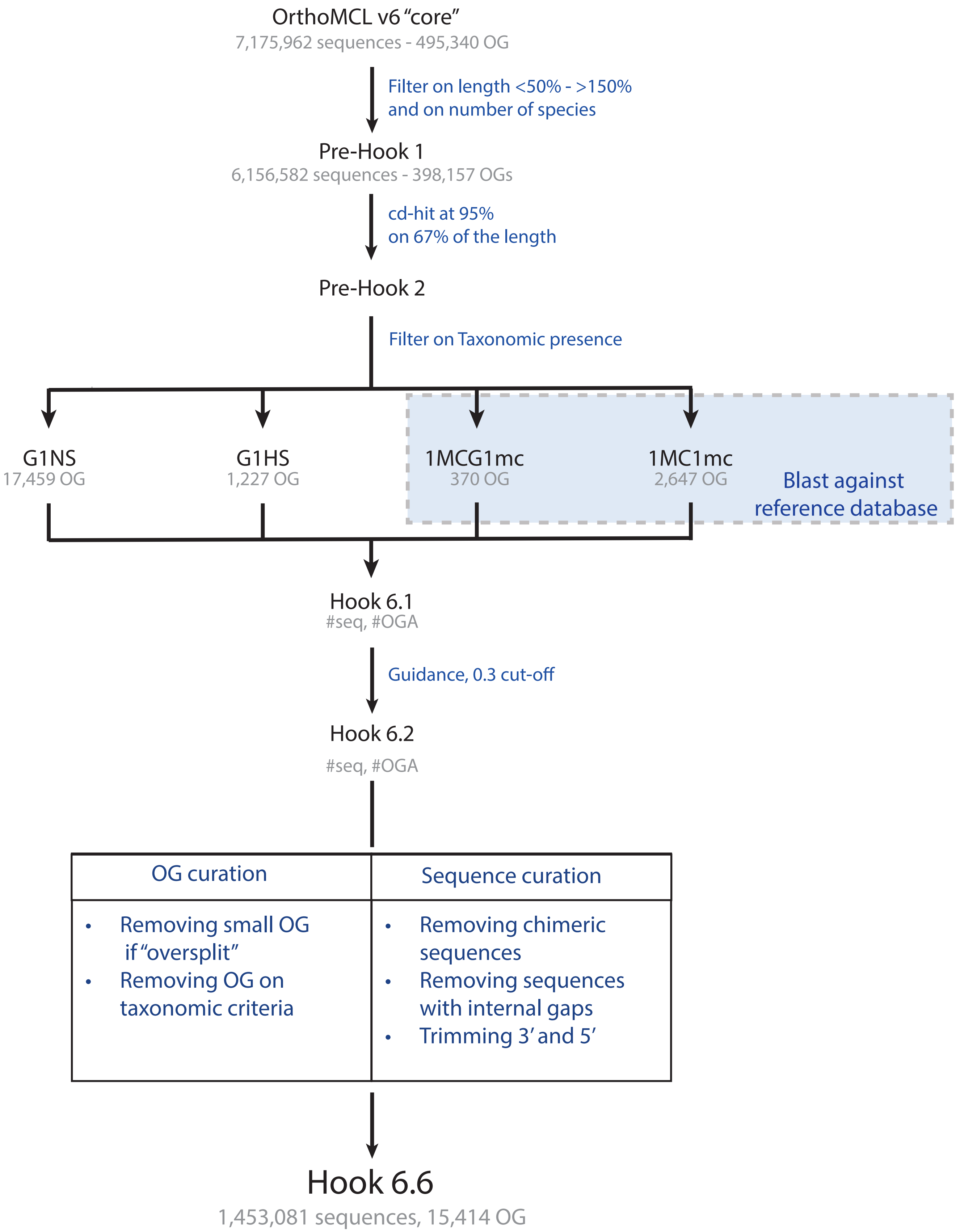

### Fig S2

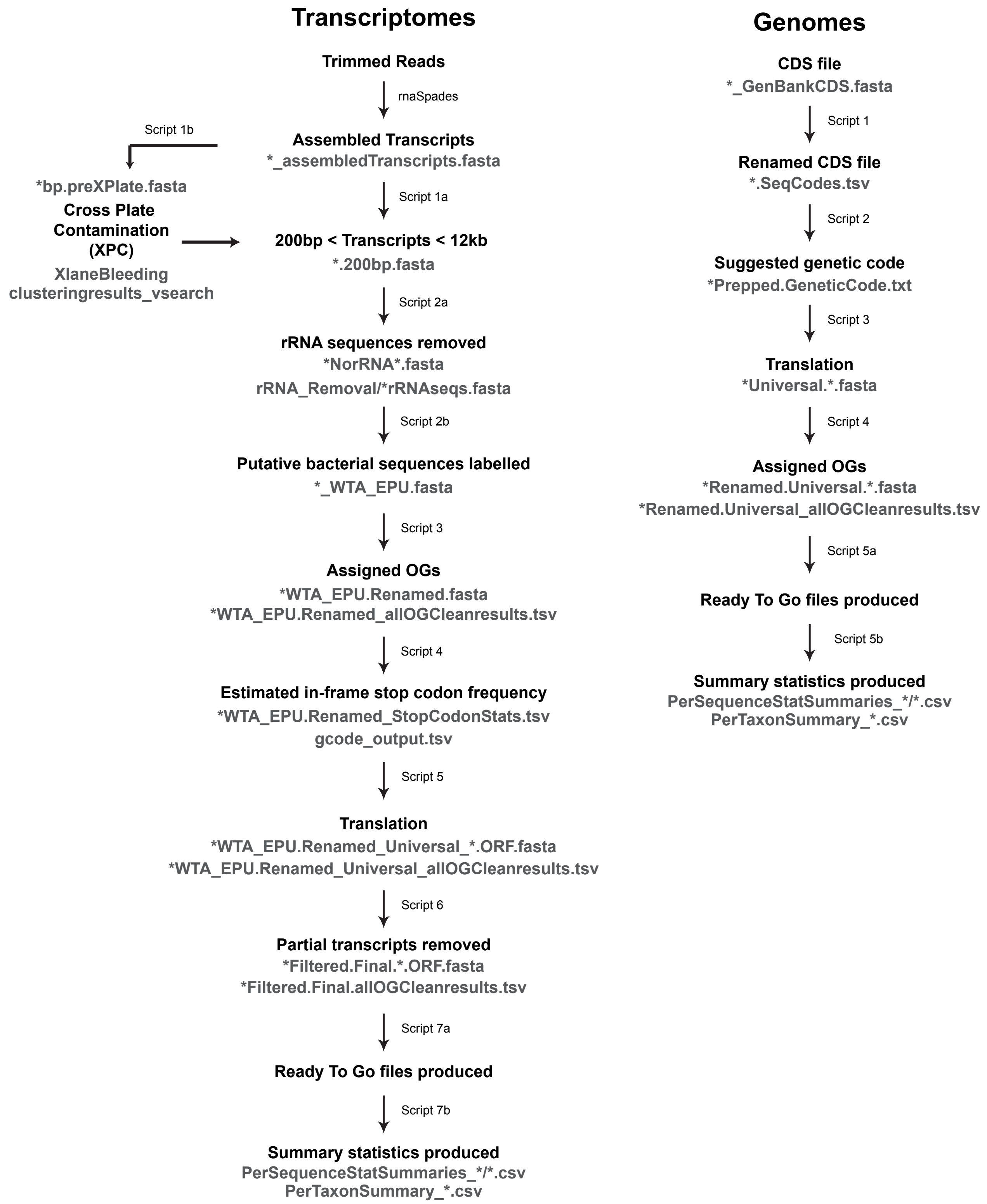

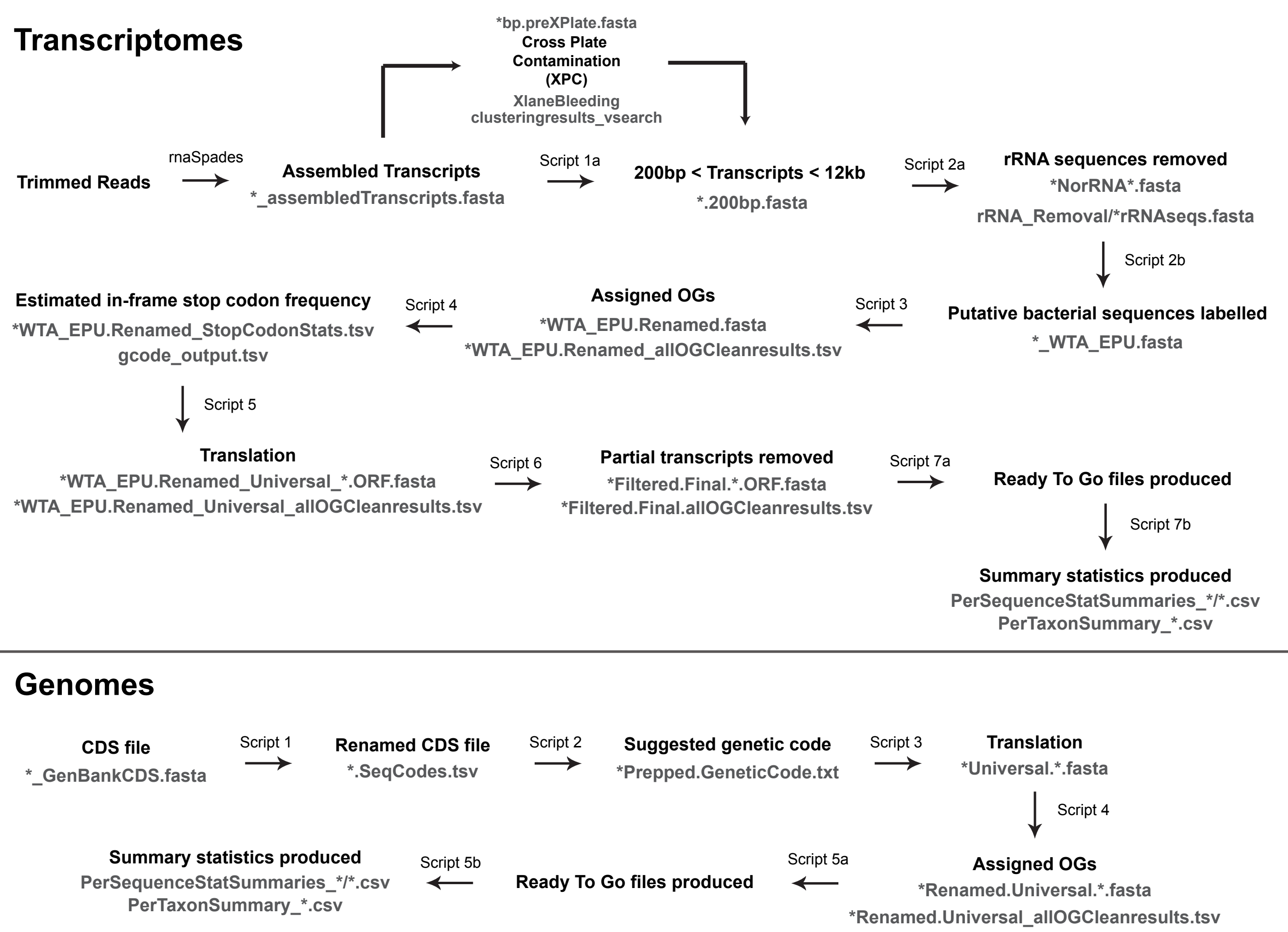

### Fig S4

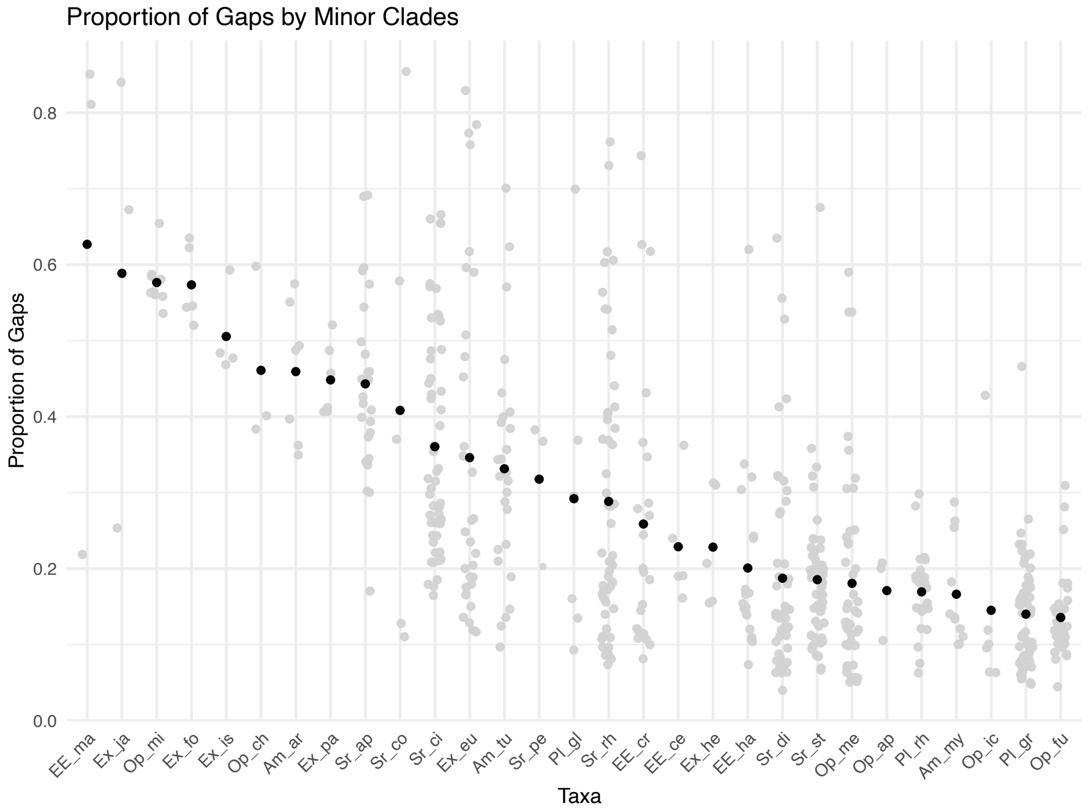

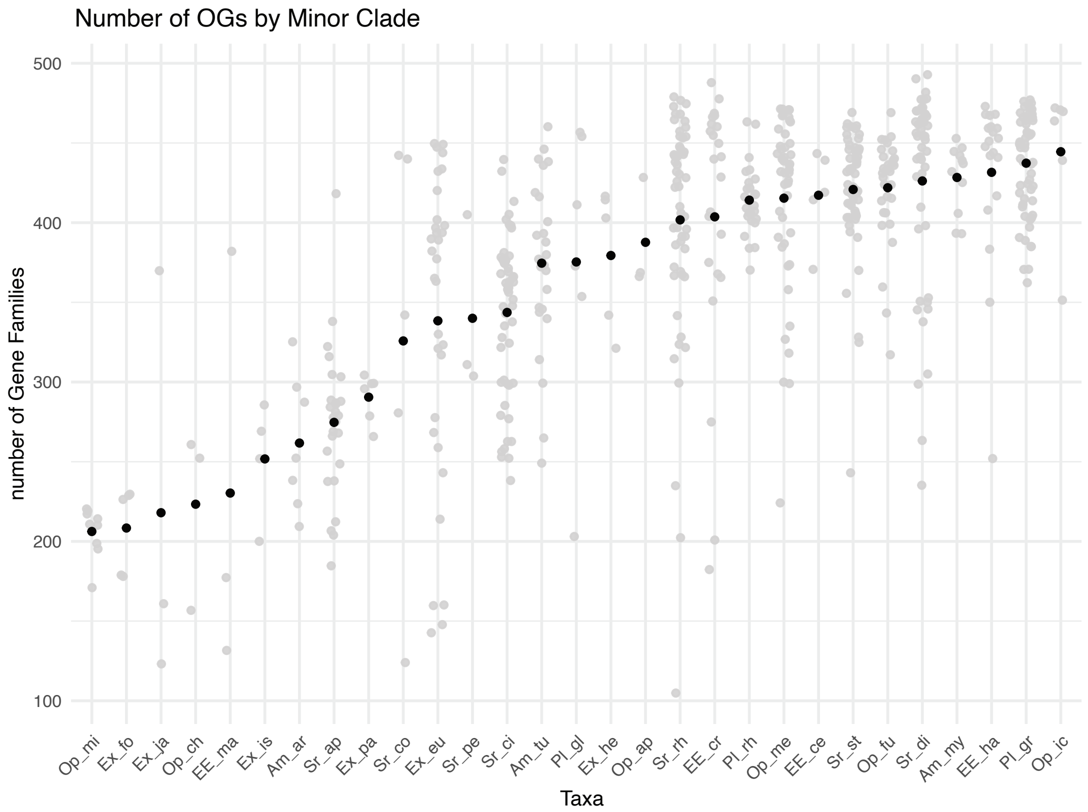
