## Supplementary material for "Rethinking large scale phylogenomics with EukPhylo v1.0, a flexible toolkit to enable phylogeny-informed data curation and analyses of diverse eukaryotic lineages": Fig S3

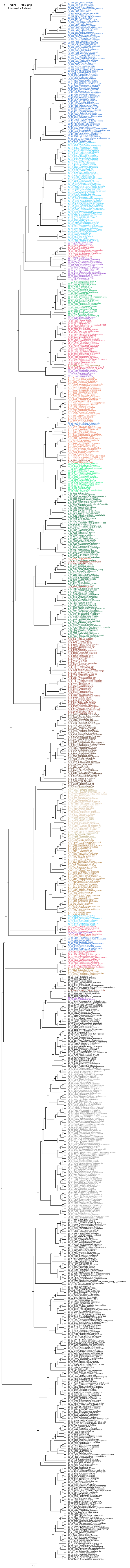
a. EndPTL - 50% gap
Trimmed - Asteroid

b. EndPTL - 50% gap Trimmed - Concatenation

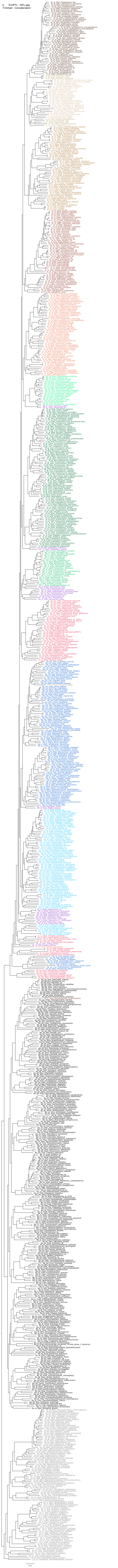

c. Sisters Removed - 50%
gap Trimmed - Asteroid

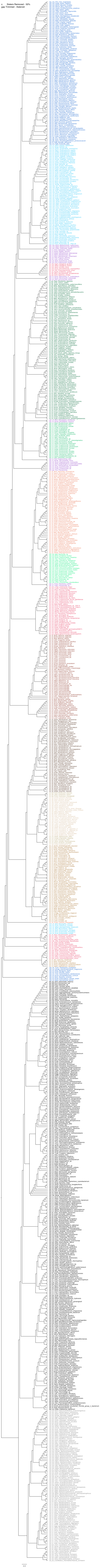

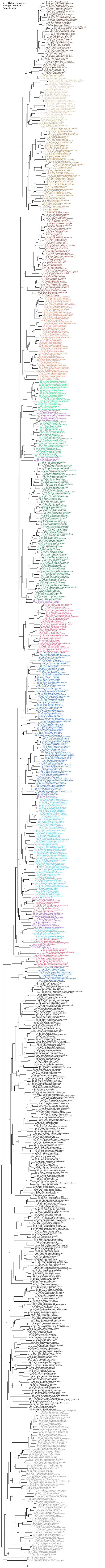
d. Sisters Removed -
50% gap Trimmed -
Concatenation
0.3

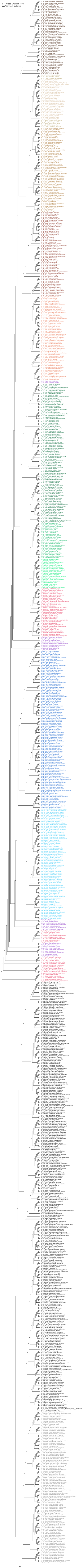
e. Clade Grabbed - 50% gap Trimmed - Asteroid
Sr_di_Pbah_Pyrodinium_bahamense
Sr_di_Gaus_Gambierdiscus_australes
Sr_di_Plun_Pyrocystis_lunula
Sr_di_Pbei_Pelagodinium_beii
Sr_di_Atam_Alexandrium_tamarense
Sr_di_Pret_Protoceratium_reticulatum
Sr_di_Gspi_Gonyaulax_spinifera
Sr_di_Cfus_Ceratium_fusus
Sr_di_Dfor_Dinophysis_fortii
Sr_di_Aspi_Azadinium_spinosum
Sr_di_Pmic_Prorocentrum_micans
Sr_di_Paci_Peridinium_aciculiferum
Sr_di_Shan_Scrippsiella_hangoei
Sr_di_Ccoh_Crypthecodinium_cohnii
Sr_di_Stro_Scrippsiella_trochoidea
Sr_di_Bnut_Brandtodinium_nutriculum
Sr_di_Kfol_Kryptoperidinium_foliaceum
Sr_di_Htri_Heterocapsa_triquetra
Sr_di_Harc_Heterocapsa_arctica
Sr_di_Sysp_Symbiodinium_sp
Sr_di_Snec_Symbiodinium_necroappetens
Sr_di_Smic_Symbiodinium_microadriaticum
Sr_di_Snat_Symbiodinium_natans
Sr_di_Spil_Symbiodinium_pilosum
Sr_di_Fkaw_Fugacium_kawagutii
Sr_di_Cgor_Cladocopium_goreaui
Sr_di_Bmin_Breviolum_minutum
Sr_di_Dtre_Durusdinium_trenchii
Sr_di_Badr_Biecheleriopsis_adriatica
Sr_di_Bcin_Biecheleria_cincta
Sr_di_Pgla_Polarella_glacialis
Sr_di_Yyeo_Yihiella_yeosuensis
Sr_di_Lfis_Levanderina_fissa
Sr_di_Tjol_Togula_jolla
Sr_di_Gcat_Gymnodinium_catenatum
Sr_di_Lchl_Lepidodinium_chlorophorum
Sr_di_Espp_Erythropsidinium_sp
Sr_di_Kbre_Karenia_brevis
Sr_di_Kmik_Karenia_mikimotoi
Sr_di_Kven_Karlodinium_veneficum
Sr_di_Amas_Amphidinium_massartii
Sr_di_Acar_Amphidinium_carterae
Sr_di_Nsci_Noctiluca_scintillans
Sr_di_Ams2_Amoebophrya_sp
Sr_di_Ams1_Amoebophrya_sp
Sr_di_Hsps_Hematodinium_sp
Sr_di_Omar_Oxyrrhis_marinia
Sr_pe_Perk_Perkinsus_marinus
Sr_pe_Olen_Perkinsus_olseni
Sr_pe_Pche_Perkinsus_chesapeaki
Sr_ap_Bbig_Babesia_bigemina
Sr_ap_Bbov_Babesia_bovis
Sr_ap_Tpar_Theileria_parva_strain_Muguga
Sr_ap_Tann_Theileria_annulata_strain_Ankara
Sr_ap_Pfal_Plasmodium_falciparum
Sr_ap_Pviv_Plasmodium_vivax
Sr_ap_Nspa_Nephromyces
Sr_ap_Tgon_Toxoplasma_gondii
Sr_ap_Hham_Hammondia_hammondi
Sr_ap_Ncan_Neospora_caninum
Sr_ap_Eace_Eimeria_acervulina
Sr_ap_Emax_Eimeria_maxima
Sr_ap_Eten_Eimeria_tenella
Sr_ap_Cpar_Cryptosporidium_parvum
Sr_ap_Chom_Cryptosporidium_hominis
Za_em_Marv_Methanocella_arvoryzae
Za_em_Mbou_Methanoculleus_bourgensis
Za_em_Mman_Methanoculleus_marisnigri
Za_em_Mlit_Methanofollis_liminatans
Za_em_Mtar_Methanolinea_tarda
Za_em_Mpal_Methanosphaerula_palustris
Za_ea_Fpla_Ferroglobus_placidus
Za_ea_Gaha_Geoglobus_ahangari
Za_ea_Aven_Archaeoglobus_veneficus
Za_ep_Acup_Acidiplasma_cupricumulans
Za_ep_Fac1_Ferroplasma_acidarmanus
Za_ep_Ptor_Picrophilus_torridus
Za_ep_Taci_Thermoplasma_acidophilum
Za_ep_Tarc_Thermoplasmatales_archaeon
Za_eu_Aboo_Aciduliprofundum_boonei
Za_eu_Aspa_Aciduliprofundum_sp
Za_em_Mlum_Methanomassiliicoccus
Za_ec_Maeo_Methanococcus_aeolicus
Za_ec_Moki_Methanothermococcus_okinawensis
Za_ec_Mign_Methanotorris_igneus
Za_ec_Mjan_Methanocaldococcus_jannaschii
Za_eb_Mbry_Methanobacterium_lacus
Za_eb_Mthe_Methanothermobacter_thermautotrophicus
Za_eb_Mrum_Methanobrevibacter_ruminantium
Za_et_Tkod_Thermococcus_kodakarensis
Za_et_Tsib_Thermococcus_sibiricus
Za_et_Pfur_Pyrococcus_furiosus
Za_cr_Msed_Metallosphaera_sedula
Za_cr_Mpru_Metallosphaera_prunae
Za_cr_Mhak_Metallosphaera_hakonensis
Za_cr_Mcup_Metallosphaera_cuprina
Za_cr_Ahos_Acidianus_hospitalis
Za_cr_Asul_Acidianus_sulfidivorans
Za_cr_Abri_Acidianus_brierleyi
Za_cr_Stok_Sulfurisphaera_tokodaii
Za_cr_Sohw_Sulfurisphaera_ohwakuensis
Za_cr_Saci_Sulfolobus_acidocaldarius
Za_cr_Sisl_Sulfolobus_islandicus
Za_cr_Ssol_Sulfolobus_solfataricus
Za_cr_Tagg_Thermosphaera_aggregans
Za_cr_Dmuc_Desulfurococcus_mucosus
Za_cr_Smar_Staphylothermus_marinus
Za_cr_Iagg_Ignisphaera_aggregans
Za_cr_Pocc_Pyrodictium_occultum
Za_cr_Hbut_Hyperthermus_butylicus
Za_cr_Pfum_Pyrolobus_fumarii
Za_cr_Ihos_Ignicoccus_hospitalis
Za_cr_Asac_Acidilobus_saccharovorans
Za_cr_Tten_Thermoproteus_tenax
Za_cr_Tuzo_Thermoproteus_uzoniensis
Za_cr_Paer_Pyrobaculum_aerophilum
Za_cr_Vdis_Vulcanisaeta_distributa
Za_cr_Vmou_Vulcanisaeta_moutnovskia
Za_cr_Cmaq_Caldivirga_maquilingensis
Za_ko_Ckor_Candidatus_Korarchaeum
Za_th_Nkor_Nitrosoarchaeum_koreensis
Za_th_Nmar_Nitrosopumilus_maritimus
Za_th_Ngar_Nitrososphaera_gargensis
Za_as_Thor_Thorarchaeota_archaeon
Za_as_Thob_Thorarchaeota_archaeon
Za_as_Odin_Odinarchaeota_archaeon
Za_as_Loka_Lokiarchaeota_archaeon
Za_as_Loki_Lokiarchaeum_sp
Za_as_Candidatus_Heimdallarchaeota_archaeon
Za_as_Heia_Heimdallarchaeota_archaeon
4.0

f. Clade Grabbed - 50% gap Trimmed - Concatenation

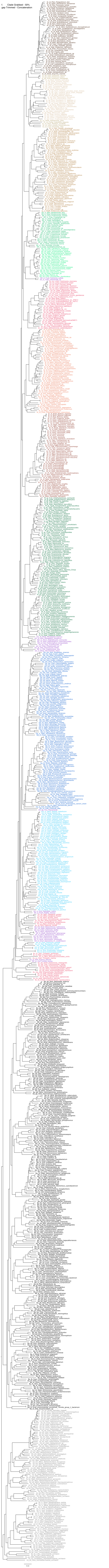

g. Putative EGT removed - 50% gap Trimmed - Asteroid

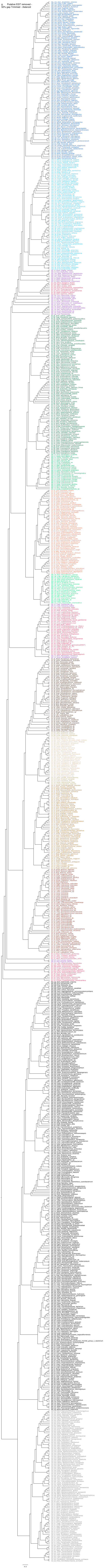

h. Putative EGT removed - 50% gap Trimmed - Concatenation

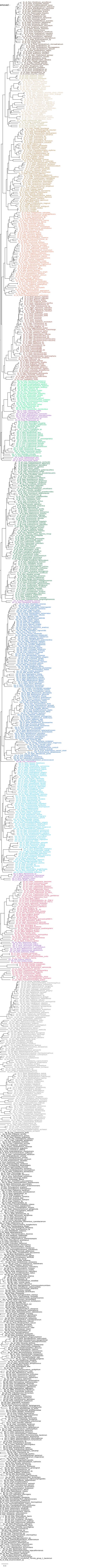
